# Phylogenetic Analysis and Machine Learning Identify Signatures of Selection and Predict Deleterious Mutations in Common Bean

**DOI:** 10.1101/2025.05.05.652309

**Authors:** Henry Cordoba-Novoa, Edward S. Buckler, Cinta M. Romay, Ana Berthel, Lynn Johnson, Parthiba Balasubramanian, Valerio Hoyos-Villegas

## Abstract

Mutations are continuous source of new alleles and genetic diversity in populations. Domestication and selection influence the accumulation of alleles occurring across a range of deleteriousness. Evidence suggests that mildly deleterious mutations (DelMut) can be purged out of breeding populations, increasing favorable allele accumulation. We used phylogeny-based analyses among 36 legume genomes to identify selection signatures and predict DelMut in common bean. We also developed a multiparent advanced generation intercrossed (MAGIC) population of black beans to characterize DelMut. Genes involved in nitrogen metabolism showed signs of positive selection in the Middle American genome, whereas genes related to phosphorylation were positively selected in the Andean genome. By combining conservation and protein information with machine learning (ML) for high-dimensional feature analysis, we characterized 82,442 sites in the MAGIC founders (36,558 polymorphic) and 4,753 sites evenly sequenced among RILs that could be potentially deleterious. Variation in the number of *highly* DelMut (high predicted deleterious scores) among lines was observed and later correlated with agronomic traits. Phenotypic analyses showed that calculated genetic load (and number of *highly* DelMut) was negatively correlated with flowering time, maturity, and yield. A detailed *in-silico* analysis of predicted mutations showed presence in highly conserved protein regions, which is likely to affect protein functionality. Our results show that variation in genetic load can be observed in breeding populations and potentially impact plant performance. These results contribute to understanding the genome-wide accumulation patterns of DelMut in breeding populations. Our study supports future development of strategies to reduce genetic load in promising germplasm and accelerate breeding programs.

**Key messages:**

- Genome-wide highly deleterious mutations were predicted in conserved protein domains potentially affecting protein functionality.
- Variation in the genetic load and number of highly deleterious mutations can be observed in artificial breeding populations.
- Nitrogen- and phosphorous-related genes are under positive selection in Middle American and Andean beans respectively.

## Introduction

Every generation, mutations are the source of new alleles and genetic diversity in populations. Depending on their effect on plant performance, mutations can be advantageous, neutral or deleterious (Roth and Liberles, 2006). Deleterious mutations (DelMut) with drastic phenotypic effects are kept at low frequency and rapidly purged from the population. On the contrary, mutations with positive effects will increase in frequency and be fixed. Due to their generally partially recessive nature, mildly DelMut escape purifying selection and tend to accumulate in populations, which is known as the genetic load (Grossen and Ramakrishnan, 2024; Kono et al., 2016).

Domestication and improvement create population bottlenecks that increase genetic drift and inbreeding (Bosse et al., 2019; Kirkpatrick and Jarne, 2000). An excess of mating of individuals closely related reduces the effective population size, which in turn reduces the efficacy of selection and increases the rate of fixation of slightly DelMut (Charlesworth, 2009; Kimura, 1983). Studies have shown that domestication is accompanied by an increase in DelMut in outcrossing and asexually propagated species compared to self-pollinated (Dwivedi et al., 2023). Because of the inbreeding dynamics, mutations with a higher effect are more easily exposed and eliminated in self-pollinated species (David et al., 1993; Wright et al., 2013). Similar to asexual populations, in the absence of recombination with mutation-free genotypes, DelMut can also accumulate in self-pollinated species through Muller’s Ratchet (Heller and Smith, 1978), a process that describes the accumulation of irreversible DelMut in the absence of recombination.

The accurate identification of DelMut and their actual effect on phenotypic variation have been a topic of extensive interest. DelMut are assumed to alter highly conserved sites and affect protein function (Yampolsky et al., 2005; Chun and Fay, 2009). Different methods based on either sequence conservation or potential effects on amino acid substitutions have been developed (Davydov et al., 2010; Ng and Henikoff, 2003; Stone and Sidow, 2005). Nevertheless, machine learning (ML) algorithms have emerged as an attractive approach for the simultaneous analysis of high-dimensional feature sets, enabling a better characterization of deleterious alleles (Adzhubei et al., 2010; Kovalev et al., 2018; Ramstein and Buckler, 2022).

Phylogenetic analyses allow the detection of selection footprints in diverse lineages (Massey et al., 2008; Yang and Wyckoff, 2011) and can be exploited for the identification of deleterious alleles (Long et al., 2023). The ratio of non-synonymous to synonymous substitutions (ω *= dN/dS*) estimates the strength and mode of natural selection on protein-coding sequences (Jeffares et al., 2015). Simultaneously, differential selective constraints influence variation in evolutionary rates across sites based on the functional and structural requirements of a gene or protein (Yang, 1996). Mutations occurring in sites with low variation rates – with proper substitution models – and predicted impact on protein function are expected to be deleterious. The study of DelMut opens new opportunities for crop improvement. As the understanding of DelMut biology and advancements of conventional and molecular breeding tools take place, purging DelMut from domesticated and wild germplasm has become an objective in crop improvement research (Gao, 2021; Wallace et al., 2018). In cotton (*Gossypium hirsutum* L.), the introgression of the *Ren^Ion^* allele for reniform nematode resistance resulted in a strong linkage drag that caused plants to be stunted, less productive and with increased susceptibility to soil-borne pathogens. In an effort to break the linkage between the nematode resistance and the stunting phenotype, Zheng et al. (2016) designed an intensive SNP-based Marker-assisted selection (MAS) with limited success. This is an example of how the identification and targeted purging of potential DelMut(s) from diverse germplasm could lead to a significant improvement of crops without yield penalties. Recently, Glaus et al. (2025) identified a missense DelMut in the tomato transcription factor *SUPRESSOR OF SP2 (SSP2)* that was highly prevalent after domestication. Using base editing, the repair of the DelMut produced more compact plants with earlier fruit production. DelMut have been predicted genome-wide in diverse maize (Mezmouk and Ross-Ibarra, 2014; Ramstein and Buckler, 2022; Yang et al., 2017), tomato (Glaus et al., 2025), sorghum (Valluru et al., 2019), barley (Kono et al., 2016), soybean (Kim et al., 2021), cassava (Long et al., 2023; Ramu et al., 2017) and potato (Wu et al., 2023) populations using different approaches, increasing the resources for breeding purposes.

Common bean (*Phaseolus vulgaris* L.) is the main legume crop for direct human consumption particularly in developing countries (Broughton et al., 2003). Common bean underwent two independent domestication events, one in Middle America (Mexico) and another in the Andes (south of northern Peru and Ecuador; Bitocchi et al., 2012). These two gene pools differ in both morphoecological and genetic characteristics, with Middle American beans exhibiting greater diversity (Schmutz et al., 2014). To date, few studies have performed genome-wide analyses of selection among common bean gene pools and no studies on deleterious mutations have been reported.

Considering the importance of crop, the highly selfing biology of the plant, and the domestication processes and diversity, we applied a phylogenetic and ML approach for the study of both selection signatures and the genomic landscape of DelMut in common bean. Our objective was to identify selection signatures in common bean genes depending on the gene pool reference genomes, and to develop a ML model that jointly leveraged conservation and protein information to predict DelMut in both Middle American and Andean beans. Once predicted, we studied the accumulation patterns of DelMut in a breeding MAGIC population of black beans (Middle American) and showed the potential protein changes induced by genome-wide identified mutations.

## Materials and methods

### Selection signatures in common bean genes

To establish the phylogenetic relationships and determine the degree of gene conservation in beans when compared to other members of the Fabaceae, we adopted an approach based on evolutionary information among species from the Fabaceae family. A total of 36 high-quality, publicly available genomes (Table S1) spanning ∼59 million years (my) of evolution and different ploidy levels were analyzed (Lavin et al., 2005). The primary transcripts of the Middle American UI111 common bean reference genome v1.1 (https://phytozome-next.jgi.doe.gov/info/PvulgarisUI111_v1_1) and the Andean G19833 v2.1 reference (Schmutz et al., 2014) were independently aligned against each legume genome using Minimap2 (Li, 2018). Introns were removed and a multiple sequence alignment (MSA) per transcript across all species was generated using MAFFT (Katoh et al., 2002). Based on the MSA for each transcript, phylogenetic trees were calculated with the Randomized Axelerated Maximum Likelihood (RAxML) method (Stamatakis, 2014; Figure 1A).

**Figure 1.**
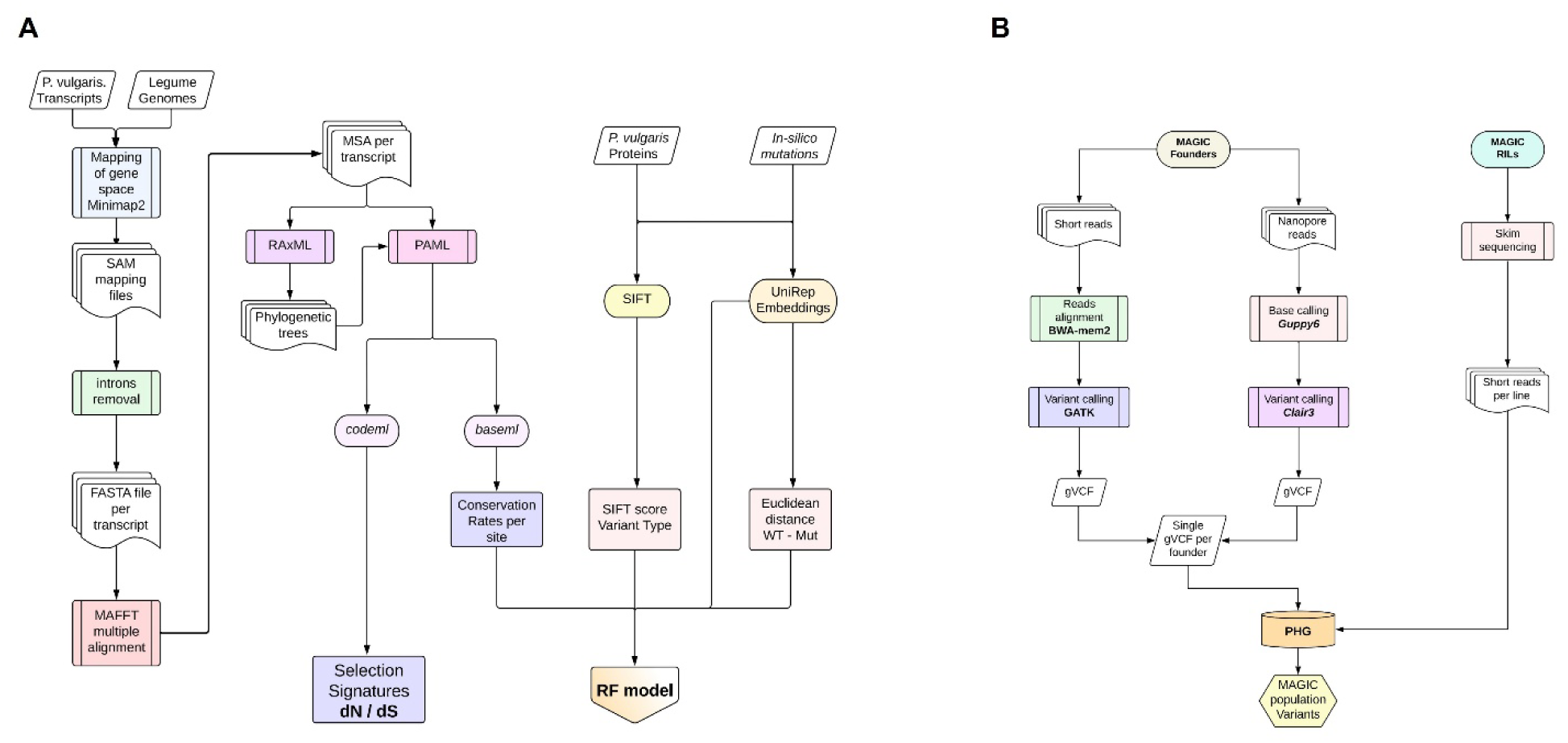
Bioinformatic pipeline for the identification of selection signatures and Random Forest (RF) model training (A), and valiant calling and imputation using the Practical Haplotype Graph (PHG) for the MAGIC population (B).

To detect selection signatures in common bean genes from both the Middle American and Andean reference genomes we used the module *codeml* from the Phylogenetic Analysis by Maximum Likelihood (PAML) suite and estimated the ratio between nonsynonymous and synonymous substitution rates (ω = *dN / dS*) according to Álvarez-Carretero et al. (2023). For each gene, two models, a null (model = 0) and a branch model (model = 2), were fitted. The null model assumes a single ω ratio for the entire phylogenetic tree, whereas the branch model allows common bean to have a different ω ratio. The models were compared based on the maximum likelihood (ML) scores with a Likelihood Ratio Test (LRT). The difference between the ratio estimated for the tree and the branch (Δ dN/dS) was used to identify differential selection in common bean compared to the other legume species. A Δ dN/dS < 0 indicates that the gene shows positive selection (more functional variation – non-synonymous mutations) in the common bean branch compared to the rest of the phylogenetic tree. A ΔdN/dS > 0 indicates a negative selection (more conservation – less variation) of the gene in common bean (Long et al., 2024). Values higher or lower than 2 and -2 were assigned 2 to facilitate visualization. Genes with significant differences between models were analyzed for Gene Ontology (GO) enrichment using AgriGO with the Fisher’s test and the reference genome annotations (Tian et al., 2017).

### Prediction of deleterious alleles

Positive selection signatures indicate that non-synonymous mutations (dN) accumulate at a higher rate than synonymous mutations (dS) in a sequence. However, some of those new variants may have a deleterious effect on the plant. To predict potential DelMut in populations, we integrated the evolutionary conservation information described above with protein function disruption data using machine learning (Long et al., 2023). Using the MSA and the phylogenetic trees previously described, variation rates were estimated for each coding base pair of the common bean genome using the module *baseml* and the General-time-reversible (GRM also known as REV) nucleotide substitution model (Yang, 1994) from PAML (Yang, 2007; Figure 1A).

Sorting Intolerant From Tolerant (SIFT) scores were calculated for the primary protein sequences of common bean (Ng and Henikoff, 2003). Briefly, SIFT analyses a reference protein database and assigns scores to non-synonymous mutations based on their likely impact on amino acid substitution and protein sequence. Lower SIFT values (close to zero) mean that the mutation will be deleterious. *In-silico* mutations were inserted in exonic sequences using a mutation rate of 1 × 10^−9^ and a divergence time of 59 million years (my) The mutation rate was defined according to previous estimates and applications in common bean (Bellucci et al., 2014; Bitocchi et al., 2017). The divergence time was based on previous phylogenetic estimates for the legume species included in this study (Lavin et al., 2005). Around 3 million variants were produced and SIFT scores were calculated for the *in-silico* mutated protein sequences. Fundamental features of normal and mutated protein sequences were summarized in protein embedding using the *256-unit* unified representation (UniRep) deep learning model (Alley et al., 2019). The Euclidian distance between the embeddings from the normal (wild type) proteins and the *in-silico* mutated was also calculated.

The following random forest (RF) model was trained with 515 features using the R package *ranger* (Wright and Ziegler, 2017) for each coding position of both the Middle American and the Andean reference genomes.

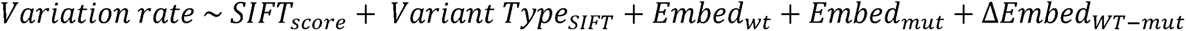

Where *SIFT* are the scores for the variants from *in-silico* mutated proteins (*mut*), *Variant type* is the category assigned by SIFT for the same variants, *Embed* is the UniRep embeddings for each kind of sequence (*wt* or *mut*), and Δ*Embed_WT-mut_* is the Euclidean difference between embeddings. The model is based on the principle that non-synonymous changes (detected within the model by SIFT and embeddings) occurring in highly conserved sites (sites with low variation rates from the MSA) are more likely to be deleterious. Once trained, the model provides a deleterious score from 0 – 1 where the higher the value the more deleterious an allele would be.

The model was trained and validated following the leave-one-chromosome-out (LOCO) approach (Ramstein and Buckler, 2022). A model was trained systematically on one chromosome at a time to predict the deleterious scores using the information from the other 10 chromosomes. For validation, sites with a variation rate < 0.5 and predictions from the RF model > 0.5 were considered conserved sites. Two independent models were trained, one suitable for the prediction of delMut in beans with Middle American backgrounds and another for genotypes with Andean backgrounds.

### Plant Material: Black bean MAGIC population

To test the trained RF model in a population that maximizes recombination (Scott et al., 2020), we developed a common bean Multiparent Advanced Generation Intercross (MAGIC) population with eight black beans from the Mesoamerican Diversity Panel (MDP) under greenhouse conditions. Based on previously reported genotypic data for MDP (Moghaddam et al., 2016), the founders parents Zorro, AC Black Diamond, AC Harblack, Black Knight, Condor, F04_2801_4_6_6, F04_2801_4_5_1, and Ui911 were *in-silico* selected based on allelic diversity maximization analysis (Ladejobi et al., 2016). For a higher number of combinations, we followed a multi-funnel or biallelic crossing scheme where all possible parental crosses were contemplated. Crosses involving the same two parents, either as female or male donors, were considered reciprocal. A total of 28 two-way crosses were mated to obtain 70 four-way crosses, later combined into 35 eight-way crosses (Figure S1). Each eight-way cross became a family, and 18 lines per family were advanced using the single seed descent (SSD) method until F_4:5_ with a final population size of 532 recombinant inbred lines (RILs).

### Field experiments and phenotyping

The MAGIC population lines were increased in the field at the Emile Lods Research Center, McGill University in Sainte-Anne-de-Bellevue, QC in 2023. Lines were planted in 5 m single-row plots with a row spacing of 0.76 m and navy (white) beans between lines to eliminate contamination. In 2024, lines were grown in an alpha-lattice design with two replicates and 108 blocks per replicate.

agronomic traits were recorded. Days to flowering (DTF) were measured as the days from planting until 50% of the plants in a plot had open flowers. Days to maturity (DTM) were recorded when around 50% of the plot reached physiological maturity and plants were at least 60% senesced. Lodging was scored at harvest on a 1 to 10 scale where 1 indicates 100% of the plants are standing erect and 10 indicates 100% of the plants are prostrated. Seed yield was calculated in Kg/Ha and corrected for seed moisture measured with a DICKEY-john^®^ GAC^™^ 2500-INTL Grain Moisture Analyzer.

### DNA sequencing

To obtain a high-quality and dense genotype dataset, the genomes of the eight founders of the population were sequenced with both long and short reads. RIL genomes were short-read sequenced only for imputation based on the parents sequencing. For long-read sequencing, seeds were germinated under dark conditions and seedlings, avoiding cotyledons, were immediately frozen in liquid nitrogen. High molecular DNA was extracted using the CTAB-based method (Doyle and Doyle, 1987), and fragments over 25 Kb were selected using the Short Read Eliminator (SRE) kit (PacBio Inc.). Libraries were prepared and sequenced using the MinION Nanopore platform with v9.4 flowcells according to the manufacturer’s specifications (Oxford Nanopore Technologies plc). For short-read sequencing, DNA was extracted from fresh leaves using the CTAB method. Libraries were prepared with the Illumina DNA PCR-Free Prep Kit following the manufacturer’s standard protocol and recommendations. Libraries were sequenced in the Illumina NovaSeq 6000 and X platforms using paired-end and 150 bp reads with a target depth of coverage of 30X for the MAGIC founders. The RILs were skim-sequenced with short reads as previously described and a minimum target depth of 3X (Novogene Corporation Inc., CA).

### Variant calling and imputation

Long reads from the parents were processed using *Guppy 6* in the *sup* mode and processed with the PvU111 reference genome using Clair3 (Zheng et al., 2022). Short-reads were mapped to the reference using BWA-mem2 (Md et al., 2019) and later processed following the GATK pipeline (McKenna et al., 2010). Genomic Variant Calling Format (gVCF) files were obtained for both methods (Clair3 and GATK) and later concatenated for each parent. Variants with missing data, quality scores lower than 30, Minor Allele Frequency (MAF) < 0.05, and coverage < 4x for the parents were filtered out before building a practical haplotype graph (PHG) database (DB) following the pipeline recommendations (Bradbury et al., 2022; Figure 1B).

For RIL variant calling, a pangenome from the eight founders in the PHG was used to map the raw reads using Minimap2. BAM alignments were used in the imputation pipeline of the PHG in homozygous mode. Variants obtained for the final 532 RILs were filtered to remove variants missing in more than 165 RILs and with a MAF < 0.05.

### Prediction of deleterious mutations in MAGIC population and genetic load

Since the MAGIC population was built with black bean market class genotypes of the Middle American gene pool, scores of variants from the RILs were predicted using the RF model trained for Middle American beans. To avoid confounding effects due to missing sites in some lines, all variants that were not genotyped in all the lines were filtered out. For a binary classification of the DelMut, the distribution and percentiles of the predicted deleterious scores were analyzed. Alleles with scores > 99% percentile were classified as *highly* deleterious either for the parents or the RILs.

To obtain a genome-wide estimate of the total genetic load per line that allowed simple comparison, we summed all the predicted scores (RF model) for each variant in each RIL depending on the presence of the derived allele according to the following equation and similar previous approaches (Wu et al., 2023).

Two indicators per line were obtained: the number of *highly* DelMut (above the percentile (threshold)) and the genetic load, which is a dimensionless number. Due to the selfing process of the RILs, and the imputation and filtering methods, only homozygous variants were considered. The genetic load per RIL was correlated with DTF, DTM, lodging and yield using Pearson’s correlation coefficient to determine its potential effects on phenotype.

### Single gene analysis of putatively deleterious alleles

We analyzed single genes based on the number of *highly* DelMut per gene (score > 0.80) and genes carrying alleles with the highest deleterious scores from the RF model. Genes with no annotation were discarded. The protein sequence from the *P. vulgaris* UI111 Mesoamerican reference genome was used for multiple sequence alignments with other legume species using ClustalW (Thompson et al., 1994). Protein domains were annotated using InterPro Scan (Jones et al., 2014). The protein sequence from the reference was also used to model the protein tertiary structure using SWISS-MODEL (Waterhouse et al., 2018) based on Alphafold2 models (Varadi et al., 2022). The impact of predicted *highly* DelMut was assessed based on the amino acid properties.

## Results

### Selection signatures in common bean genes

The analysis of ω branch models between common bean reference genomes and other Fabaceae species showed that 1,405 genes with Middle American background and 1,231 genes with Andean background had significant selection signatures (positive or negative). In the Middle American genome, 980 genes were under positive selection (Δ *dN/dS* < 0) and 425 under negative selection (Δ *dN/dS* > 0; Figure 2A; Table S2). In the Andean G19833 reference, 913 genes showed signatures of positive selection and 318 of conservation (Figure 2B; Table S3).

**Figure 2.**
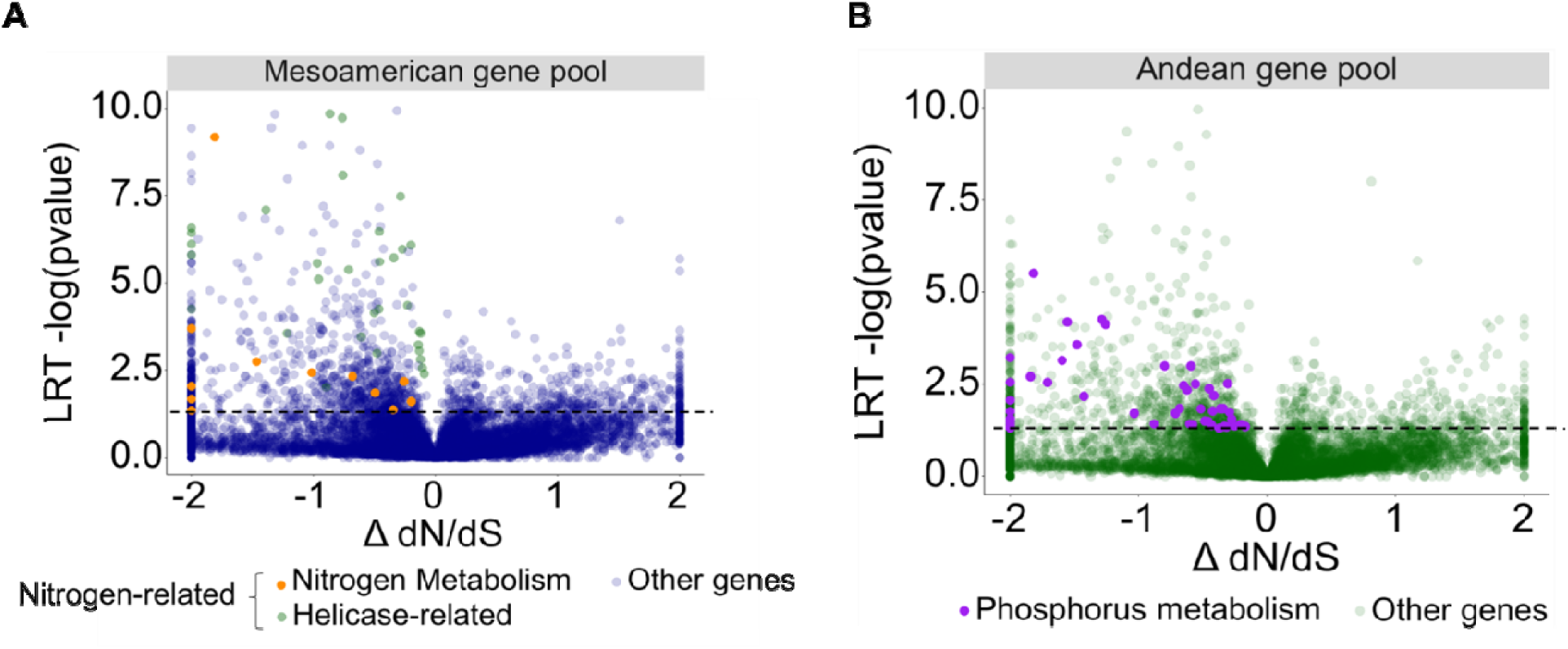
Selection signatures expressed by Δ dN/dS for genes in the Mesoamerican UI111 (A) and Andean G19833 reference genomes (B) under branch and whole tree PAML models. The dashed horizontal line denotes the threshold (*p < 0.05*) for significant differences.

In the GO enrichment analysis for the significant genes under positive selection, 25 biological processes and one Molecular Function were significant (*p<* 0.0001) in Middle American beans. We identified terms related to telomere organization and maintenance, anatomical structure, and general metabolic processes (Table 1). Unlike most terms that described generic and major metabolic pathways, two significant terms were nitrogen-related. The terms GO:0034641 and GO:0006807, had 42 genes annotated as PIF1-like helicase (PIF1) and 12 directly involved in nitrogen metabolism (Figure 2A; Table 1). For the Andean genome, four biological processes and two molecular functions were significantly enriched (*p* < 0.02) in genes undergoing positive selection. The significant terms GO:0006468 (*p* = 0.00056) – Protein amino acid Phosphorylation and GO:0016310 (*p* = 0.019) – Phosphorylation had 42 common genes related to kinase activities (Table 2). No significant GO terms were identified for genes under negative selection in the Middle American or Andean genomes.

**Table 1.**
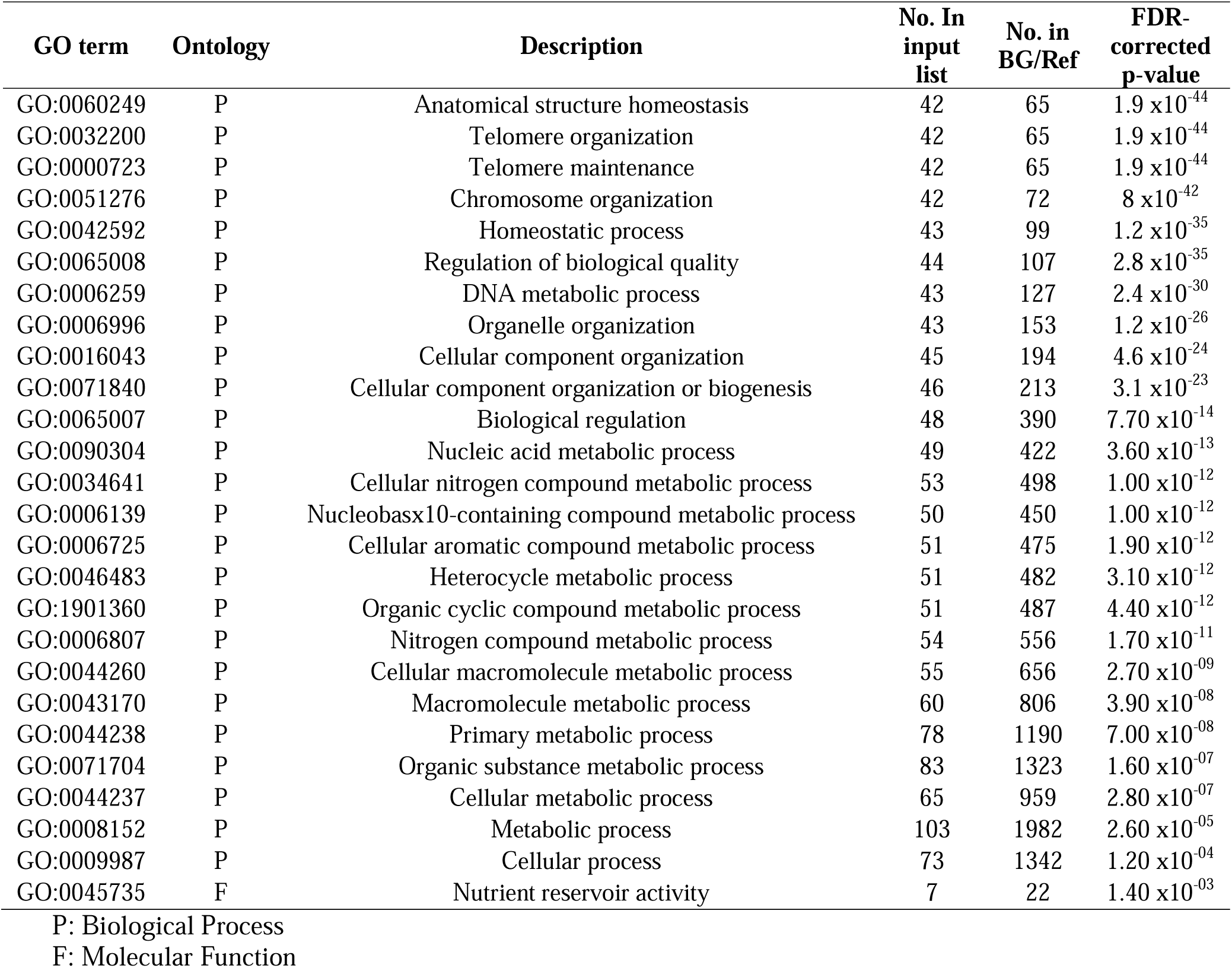
Gene Ontology (GO) Enrichment analysis for significative genes under positive selection in the Mesoamerican bean UI111 genome.

| GO term | Ontology | Description | No. In<br>input<br>list | No. in<br>BG/Ref | FDR-corrected<br>p-value |
| --- | --- | --- | --- | --- | --- |
| GO:0060249 | P | Anatomical structure homeostasis | 42 | 65 | $1.9 \times 10^{-44}$ |
| GO:0032200 | P | Telomere organization | 42 | 65 | $1.9 \times 10^{-44}$ |
| GO:0000723 | P | Telomere maintenance | 42 | 65 | $1.9 \times 10^{-44}$ |
| GO:0051276 | P | Chromosome organization | 42 | 72 | $8 \times 10^{-42}$ |
| GO:0042592 | P | Homeostatic process | 43 | 99 | $1.2 \times 10^{-35}$ |
| GO:0065008 | P | Regulation of biological quality | 44 | 107 | $2.8 \times 10^{-35}$ |
| GO:0006259 | P | DNA metabolic process | 43 | 127 | $2.4 \times 10^{-30}$ |
| GO:0006996 | P | Organelle organization | 43 | 153 | $1.2 \times 10^{-26}$ |
| GO:0016043 | P | Cellular component organization | 45 | 194 | $4.6 \times 10^{-24}$ |
| GO:0071840 | P | Cellular component organization or biogenesis | 46 | 213 | $3.1 \times 10^{-23}$ |
| GO:0065007 | P | Biological regulation | 48 | 390 | $7.70 \times 10^{-14}$ |
| GO:0090304 | P | Nucleic acid metabolic process | 49 | 422 | $3.60 \times 10^{-13}$ |
| GO:0034641 | P | Cellular nitrogen compound metabolic process | 53 | 498 | $1.00 \times 10^{-12}$ |
| GO:0006139 | P | Nucleobasx10-containing compound metabolic process | 50 | 450 | $1.00 \times 10^{-12}$ |
| GO:0006725 | P | Cellular aromatic compound metabolic process | 51 | 475 | $1.90 \times 10^{-12}$ |
| GO:0046483 | P | Heterocycle metabolic process | 51 | 482 | $3.10 \times 10^{-12}$ |
| GO:1901360 | P | Organic cyclic compound metabolic process | 51 | 487 | $4.40 \times 10^{-12}$ |
| GO:0006807 | P | Nitrogen compound metabolic process | 54 | 556 | $1.70 \times 10^{-11}$ |
| GO:0044260 | P | Cellular macromolecule metabolic process | 55 | 656 | $2.70 \times 10^{-09}$ |
| GO:0043170 | P | Macromolecule metabolic process | 60 | 806 | $3.90 \times 10^{-08}$ |
| GO:0044238 | P | Primary metabolic process | 78 | 1190 | $7.00 \times 10^{-08}$ |
| GO:0071704 | P | Organic substance metabolic process | 83 | 1323 | $1.60 \times 10^{-07}$ |
| GO:0044237 | P | Cellular metabolic process | 65 | 959 | $2.80 \times 10^{-07}$ |
| GO:0008152 | P | Metabolic process | 103 | 1982 | $2.60 \times 10^{-05}$ |
| GO:0009987 | P | Cellular process | 73 | 1342 | $1.20 \times 10^{-04}$ |
| GO:0045735 | F | Nutrient reservoir activity | 7 | 22 | $1.40 \times 10^{-03}$ |

P: Biological Process
F: Molecular Function

**Table 2.** Gene Ontology (GO) Enrichment analysis for significative genes under positive selection in the Andean G19833 genome.

| GO term | Ontology | Description | Number<br>in input<br>list | Number in<br>BG/Ref | FDR-corrected<br>p-value |
| --- | --- | --- | --- | --- | --- |
| GO:0055114 | P | Oxidation reduction | 60 | 1364 | 1.8 x10<sup>-05</sup> |
| GO:0006468 | P | Protein amino acid phosphorylation | 48 | 1134 | 5.6 x10<sup>-04</sup> |
| GO:0055085 | P | Transmembrane transport | 30 | 648 | 5.6 x10<sup>-03</sup> |
| GO:0016310 | P | Phosphorylation | 48 | 1354 | 1.9 x10<sup>-02</sup> |
| GO:0043531 | F | ADP binding | 18 | 167 | 7.8 x10<sup>-06</sup> |
| GO:0005515 | F | Protein binding | 84 | 2709 | 1.1 x10<sup>-02</sup> |

P: Biological Process
F: Molecular Function

### Model training for the prediction of deleterious mutations

For each position of the coding regions of the common bean genome, variation rates (conservation) and coverage (number of species where the position is aligned) were calculated. Rates varied from 0 to 2 and coverage from 1 to 36 (total number of species). We observed an enrichment of sites with variation rates lower than 0.5 and high coverage among the analyzed genomes (Figure 3A).

**Figure 3.**
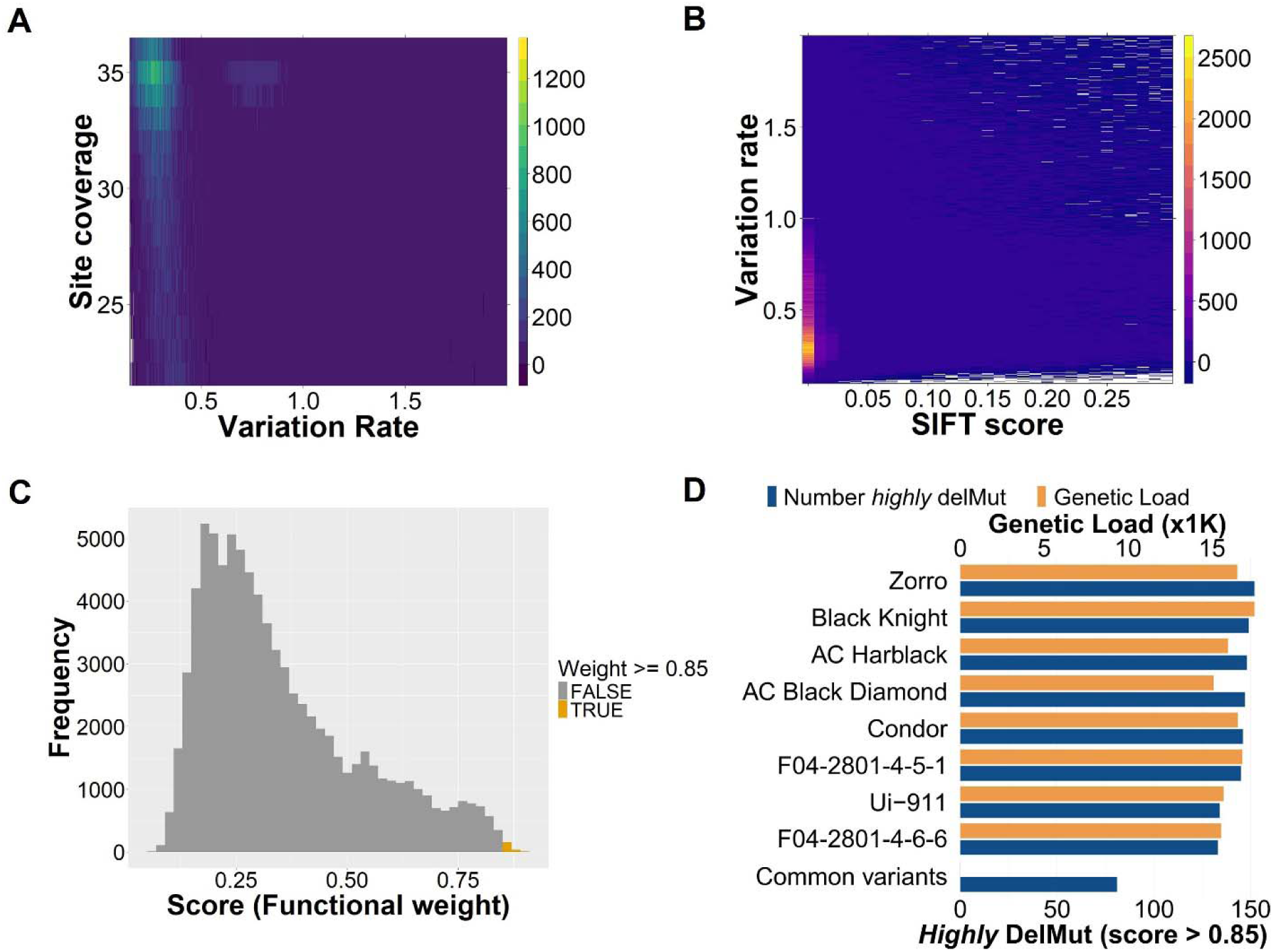
Relation between variation rate and site coverage (A) and variation rate and SIFT score (B) for RF training. Distribution of the predicted functional weights (deleterious score) from the RF model in the MAGIC population founders (C). Number of *highly* deleterious alleles (99.8% percentile) and genetic load per founder (D).

Since the model is intended to combine conservation with protein information, we analyzed the individual relationship between SIFT scores and variation rates. The estimated variation rates were correlated positively correlated with the predicted SIFT scores for each site. Sites where SIFT predicts a high impact (Low SIFT score) corresponded to sites with a low variation rate (conserved; Figure 3B). Based on the distribution of both the rates and coverage, 414,933 sites with rates lower than 0.5 and coverages greater than 21 (median) were defined as conserved and used for model training. This allows to focus the analyses and predictions on the truly conserved sites among legume species. Based on the LOCO approach, the mean accuracy of the model across chromosomes was 0.72 and the precision was 0.74, predicting high deleterious scores or functional weights for conserved sites.

### Variant calling and annotation of variants from the MAGIC population

From the processing using the PHG database, 2.7M variants were obtained among the MAGIC founders after QC filtering. In the parents, 82,443 positions had predicted scores with 53,049 genotyped in all eight founders (no missing calls). Out of these variants, 36,558 were polymorphic and 16,491 had the same allele (reference or alternative) in all the parents. As expected, the distribution of the scores (functional weights) was right skewed with an enrichment of weights from 0 to 0.5 (Figure 3C). Depending on the percentile considered for the weights, we observed variation in the number of *highly* DelMut among parents. With a more conservative threshold of 0.85 (99.8% percentile), the number of homozygous *highly* DelMut per parent varied from 133 in F04_2801_4_6_6 to 152 in Zorro with a mean of 144 DelMut (Figure 3D). Interestingly, 81 *highly* DelMut were identified as common in all the parents. Regarding the genetic load, parents had loads from 151K in Black Diamond to 17.5K in Black Knight with a mean of 14.4K (Figure 3D). For the RILs, 1.5M variants were imputed from the PHG for the whole genome. A total of 39,433 positions (2.6%) had a predicted score. Due to sequencing variability among lines where some sites were missing for some lines, a final set of 4,753 variants (12% of those with score) with predicted scores were fully genotyped in all the RILs. With a threshold of 0.80 (99% percentile), the number of *highly* DelMut (score > 0.80) per chromosome varied from 11 on chromosome 9 to 57 on chromosome 10 (Figure 4A). Variants with scores were distributed along the chromosomes with some high score SNPs (more deleterious) being common in pericentrometric regions. However, variants with high scores were also observed at the far ends of the chromosomes (Figure 4B). A similar distribution of scores was observed in the parents with a difference in the density attributable to a higher sequencing depth (Figure S2). The number of *highly* DelMut per line ranged from zero to 19 with a mean of nine (Figure 4C). The genetic load per line varied from 196 to 635 (Figure 4D) with an average value of 353 and was positively correlated (R = 0.29 – 0.73) with the binary classification when high thresholds (0.7 – 0.8) were considered (Figure S3).

**Figure 4.**
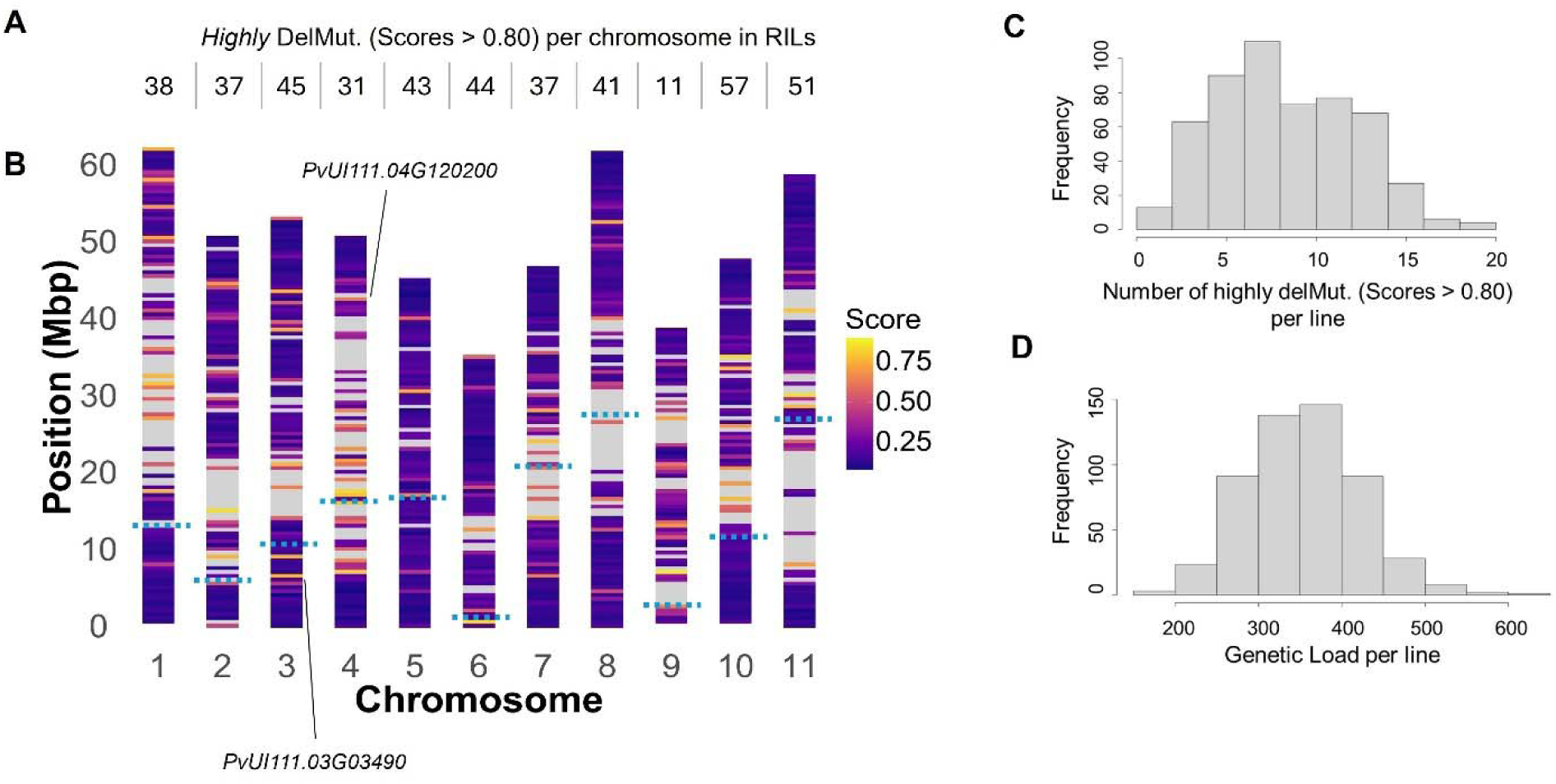
Number (A) and distribution with two high DelMut scoring genes indicated (B) of highly deleterious mutations (score > 0.80) per chromosome in the RILs. Number of highly DelMut (C) and genetic load (D) per line in the MAGIC population of black bean. Horizontal dashed blue lines indicate the telomeres according to Schmutz et al. (2014).

### MAGIC population phenotypic evaluation and correlation with genetic load

In the evaluation of agronomic traits, flowering varied from 41 to 54 days with a mean of 46 days and a SD of 2.18. Maturity ranged from 78 to 118 days with a mean value of 86 days. In lodging, lines with upright architecture (score 1) and almost fully prostrate growth habit (score 9) were observed with a mean of 4. Yield widely varied from 66.59 Kg/Ha to a maximum of 3,547.82 Kg/Ha with a mean of 1,877 Kg/Ha and a CV of 26% (Table 3; Figure 5A-D).

**Figure 5.**
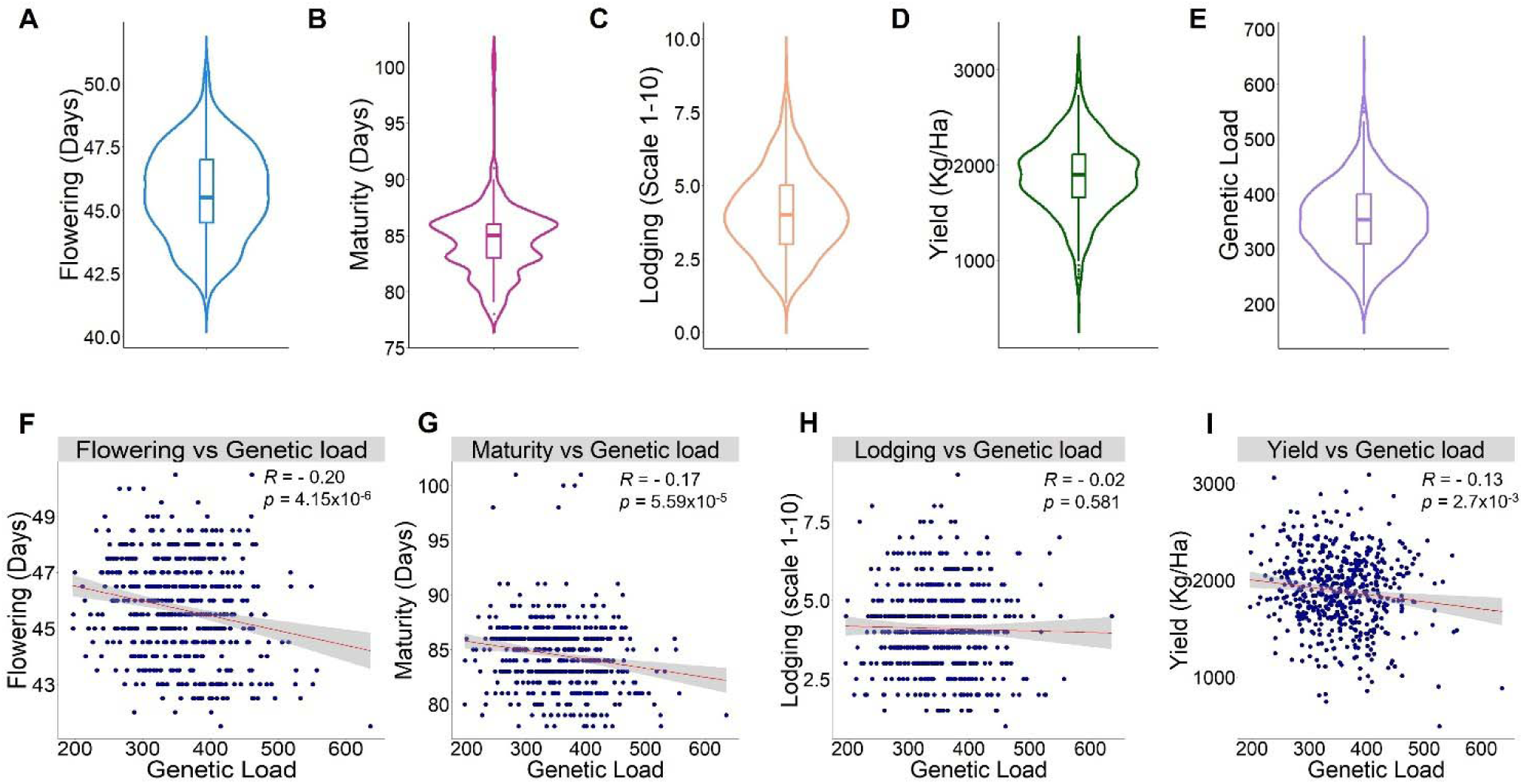
Phenotypic distribution of agronomic traits evaluated in the MAGIC population (A-D) and of the calculated genetic load per line (E). Correlation between the genetic load and the evaluated traits (F-I). Pearson’s correlation coefficient – R is indicated for each plot with the respective p-value for significance.

**Table 3.**
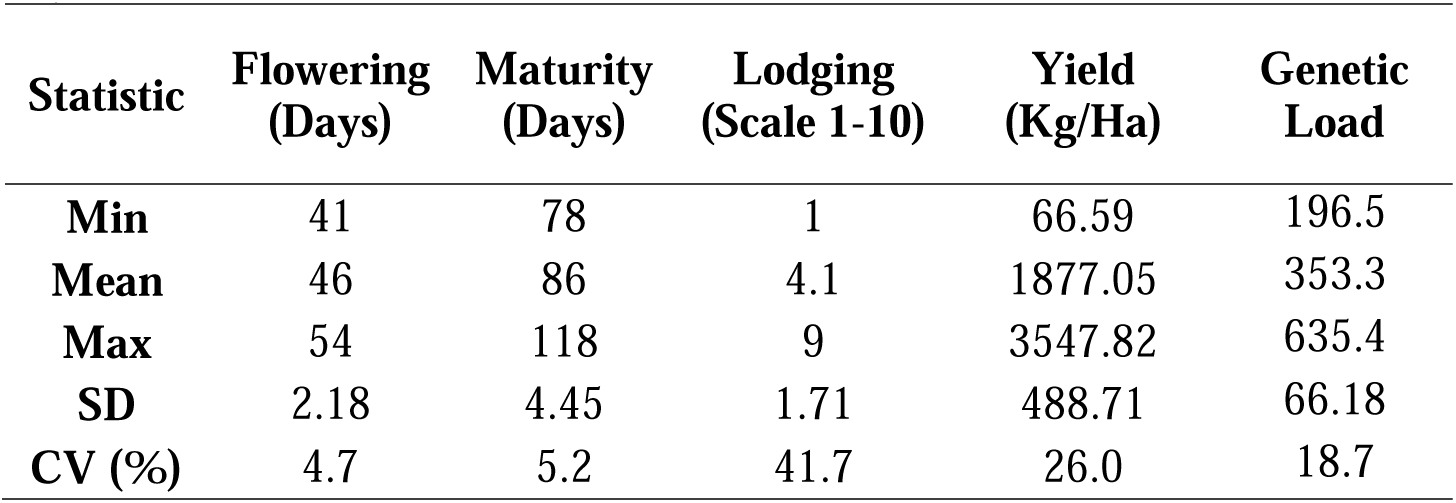
Phenotypic field evaluation of the MAGIC black bean population.

The genetic load in the RILs varied from 196.5 to 635.4 with a mean of 353.34 a SD of 66.19 and a CV of 18% (Table 3; Figure 4D and 5E). According to he Pearson correlation coefficients, flowering, maturity and yield were significantly negatively correlated (*p* < 1×10^−4^ to 0.01) with genetic load per line (R= -0.20 to -0.13). No significant correlations were detected between lodging and the genetic load (Figure 5F-I).

### Single gene analysis of putatively deleterious alleles

Among RILs, 322 gene models were predicted to accumulate between 1 – 3 *highly* (score > 80) DelMut (Table S4). We analyzed some examples of genes with the highest number of predicted DelMut and the gene with the DelMut with the highest score. The gene model *PvUI111.03G034900* coding for a Dihydrolipoyllysine-residue acetyltransferase accumulated three putatively deleterious alleles (scores = 0.82). The SNP S03_3610126 [C/T] causes the substitution of the Serine 15 residue for a Leucine (S15L) in the Biotinyl/lipoyl domain of the protein (Figure 6). Interestingly, this gene belongs to the GO term 0008152 – Metabolic process, which was significant in the GO enrichment analysis of selection signatures for the Mesoamerican genome (Figure 2A; Table 1).

**Figure 6.**
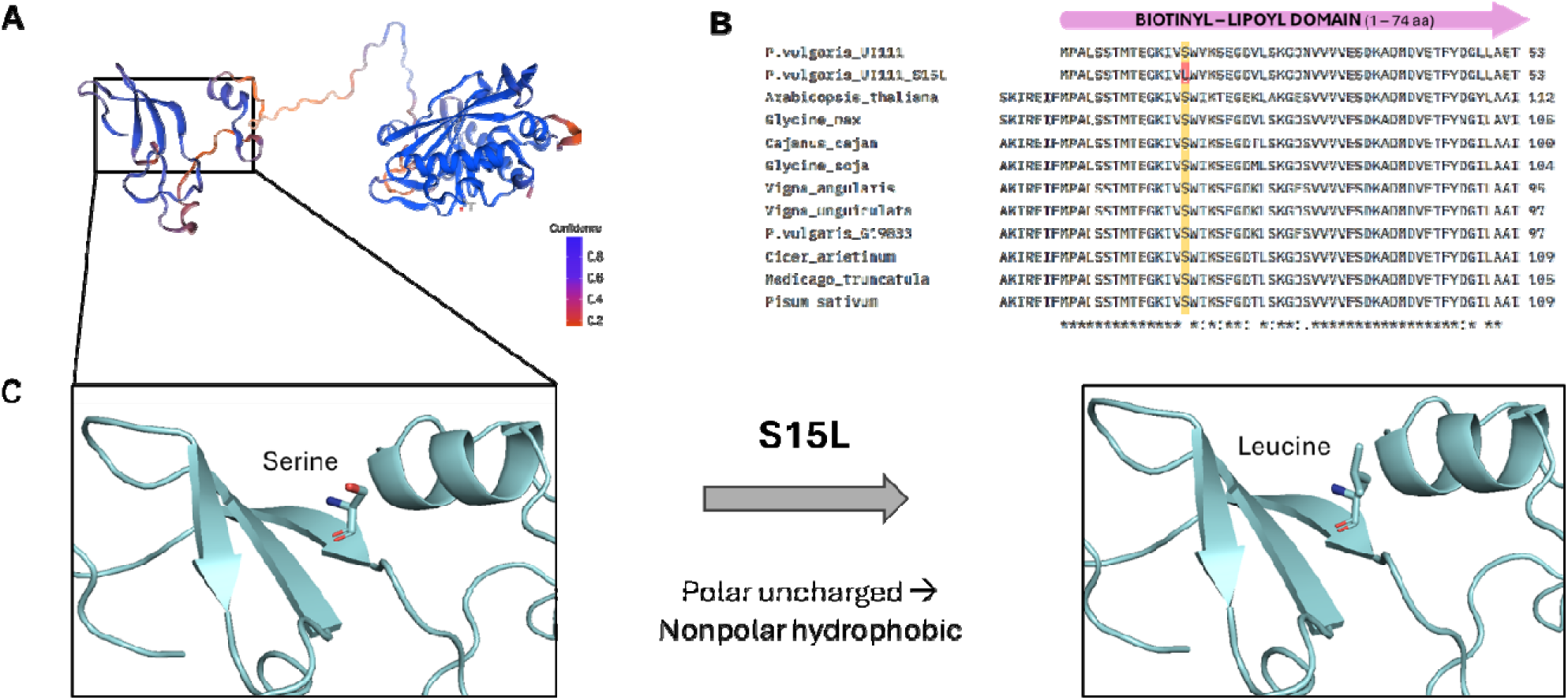
Predicted tertiary structure (A) and multiple sequence alignment (B) of the *P. vulgari* UI111 Dihydrolipoyllysine-residue acetyltransferase protein (*PvUI111.03G034900*). Amino acid substitution predicted as deleterious in the Biotinyl-Lipoyl protein domain (C).

The SNP S04_43764780 [C/G] had the highest deleterious score (0.90) and was in the coding region of the gene model *PvUI111.04G120200* – Carboxylesterase 9-related protein. The SNP causes the substitution of the Arginine 174 for Glycine (R174G) in the annotated ABHydrolase-3 domain (ABHD3; Figure S4).

## Discussion

### Selection signatures in Middle American and Andean genomes

The expansion of genomic and computational resources has increased our ability to study and leverage evolutionary information. Phylogenetic analyses have been used for the detection of selective pressures in humans, mammals, and plants (Feng et al., 2024; Massey et al., 2008; Roth and Liberles, 2006). In common bean, domestication and breeding resulted in gene loss due to selection (Cortinovis et al., 2024). In Mesoamerican genotypes, domestication also affected the levels of expression, being attributed to the presence of loss-of-function mutations (Bellucci et al., 2014). Studying lima (*Phaseolus lunatus* L.) and common bean, Li et al. (2015) detected selection footprints in genes involved in photosynthetic activity and resistance to biotic and abiotic stresses. This highlights the versatility of the *Phaseolus* genome favoring functional variation and even gene loss in different groups depending on where the gene pools have evolved.

We identified greater functional variation (more non-synonymous mutations) in nitrogen-related genes in the Mesoamerican genome and in phosphorylation-related genes in the Andean genome compared to the other analyzed legumes. Despite a few studies comparing both gene pools (Ramaekers et al., 2013; Wilker et al., 2019), Mesoamerican beans have been identified to have a higher nitrogen fixation capacity compared to Andean beans (Wilker et al., 2019). The hypothesis has been that genes controlling nitrogen fixation differ among gene pools and that greater genetic diversity in Mesoamerican beans plays a paramount role (Schmutz et al., 2014; Wilker et al., 2019). Our phylogenetic analyses support previous findings and hypothesis, indicating that N-related genes were subject to positive selection in Mesoamerican beans. Differences with the Andean genome may be partially explained by the evolutionary environment. The Andean gene pool is primarily distributed across the Andes, south of Colombia (Bitocchi et al., 2012; Kami et al., 1995). The Andisols, Oxisols and Ultisols in South America are characterized for having a low content of available phosphorous (Chacón et al., 2015; Darela-Filho et al., 2024). This may explain the prioritization of functional variation in genes related to phosphorylation in the Andean genome rather than N-related compared to other legumes. Additional analyses of specific genes under selection on diverse Andean backgrounds are needed.

### Prediction of deleterious mutations using conservation, protein information, and machine learning

Domestication, selection (natural or artificial), and mating systems affect the accumulation of deleterious alleles in populations (Bosse et al., 2019; Glémin et al., 2003; Mezmouk and Ross-Ibarra, 2014). A new allele or mutation may be deleterious under wild conditions and subject to negative (purifying) selection, yet beneficial for breeding purposes and under positive selection (Dwivedi et al., 2023). Consequently, the annotation of deleterious alleles has been mainly based on sequence conservation approaches such as GERP++ (Davydov et al., 2010) and SIFT (Ng and Henikoff, 2003) in different contexts. Depending on the set thresholds, these approaches provide binary classifications of DelMut in populations. In GERP, codon models, sequence turnovers, and missing data limit its use. Moreover, GERP scores may not accurately predict the strength of purifying selection at individual sites as both weakly and strongly DelMut can be assigned high values (Huber et al., 2020). SIFT is based on protein sequences and uses median conservation scores that may cause oversight of DelMut in highly conserved areas (Chun and Fay, 2009). By combining phylogenetic and sequence conservation information with machine-learning, we annotated alleles on a continuous scale (0 – 1). This allows a better understanding of how impactful (deleterious) an allele is predicted to be. In addition, thresholds to define subcategories (e.g. *highly* deleterious) can be easily established to prioritize certain mutations based on the population development and germplasm characteristics.

Studies in multiple crop species have shown different patterns of accumulation of DelMut depending on the reproduction system. Clonally propagated species have accumulated considerable amounts of DelMut masked in heterozygous states due to the lack of recombination (Ramu et al., 2017; Wu et al., 2023). After domestication, outcrossing species like maize tend to accumulate more deleterious mutations compared to their wild relatives, attributed to the linkage between beneficial and deleterious alleles (Wang et al., 2017). However, DelMut are later purged in elite varieties (Yang et al., 2017). For self-pollinating species like sorghum and soybean, landraces and improved varieties have less deleterious alleles than wild accessions (Kim et al., 2021; Valluru et al., 2019). As inbreeding increases, highly deleterious alleles are exposed and the purifying selection purges them out of the population. Nevertheless, mildly recessive deleterious alleles are accumulated at higher rates (Charlesworth and Willis, 2009; Fox et al., 2008). Similar to our results, Mezmouk and Ross-Ibarra (2014) and Kono et al. (2016) observed in maize and other crops, close to uniform distribution of DelMut across chromosomes, with some mutations being frequent in pericentromeric regions where recombination rate is expected to be lower. The founders of our MAGIC population are improved genotypes (Moghaddam et al., 2016). Therefore, our results are mainly limited to the accumulation of DelMut within a practical breeding context in artificial populations. Additional studies involving wild relatives, landraces, and close species within *Phaseolus* would provide new insights into the patterns of accumulation of DelMut during bean domestication and improvement. The five domesticated *Phaseolus* species may provide a platform for the study of DelMut in the context of varying mating systems. For instance, the runner bean (*P. coccineus* L.) is characterized by a high outcrossing rate compared to the other *Phaseolus* species (González et al., 2014).

Few studies have analyzed the distribution of DelMut in artificial populations in the context of breeding, with no reports for common bean. The study of DelMut is crop-specific and the number can vary from one model to another depending on the genotyping quality and approach to predict DelMut. However, variation in the number of *highly* DelMut and the genetic load among genotypes has been observed in maize and potato (Sun et al., 2023; Wu et al., 2023) where it is possible to observe in some cases close-to-normal distributions. The identification of DelMut is dependent on sequencing depth of coverage and quality. We applied a strict filter where variants that were not present in 100% of the RILs (530) were filtered out. Depending on the sequence quality and imputation methods, more variants can be predicted per population. The observed presence of variation among parents and RILs demonstrates that complex crossing schemes like the one from a MAGIC population contribute to the purging of deleterious mutations in breeding lines. Multiparent populations capture increased genetic recombination and genetic variation (Scott et al., 2020). Meiotic recombination is the main driver for the purging of deleterious mutations and increased fitness in populations (Epstein et al., 2023; Muller, 1964). In practice, the annotation of DelMut in breeding materials may contribute to the selection of parents and the design of crossing schemes to try purge them out of the breeding population and minimize their effect.

Computational biology, machine learning, and artificial intelligence (AI) methods for plant biology are rapidly evolving fields. New tools with potential applications for the prediction of DelMut in crop species have emerged in the last few years. Like UniRep, other representation and modelling of protein features for ML and AI tools have been developed. Rives et al. (2020) represented 250 million protein sequences in an unsupervised model with an Evolutionary Scale Modeling (ESM) approach. Elnaggar et al. (2022) developed ProtTrans using Language Model (LM) for protein embeddings. In the DNA space, Zhai et al. (2024, *preprint*) developed a LM for cross-species modelling and the identification of causal variants and DelMut with outstanding examples in maize. Here, we combined fundamental phylogenetic analyses with previously adopted tools for the prediction of DelMut in common bean. As novel tools and models are developed, new studies aimed at the characterization of causal and deleterious variants in common bean and other species will increase, facilitating crop improvement.*Potential effect of deleterious mutations on plant phenotypes*

The effect of the DelMut variation on plant fitness and productivity is not completely understood (Kovalev et al., 2018). Due to the recessive nature of mildly deleterious alleles, phenotypic effects, and clear, strong correlations are difficult to observe. Weak but significant negative correlations, such as those we observed between DelMut (and genetic load) and agronomic traits, have been reported. Yang et al. (2017) found that the number of homozygous SNPs classified as deleterious in maize hybrids was negatively correlated with plant height, ear height, and grain yield. More recently, Wu et al. (2023) observed in a F_2_ hybrid potato population (n = 1,064) that the homozygous burden was also negatively correlated with yield (R = -0.33), plant height (R = -0.25), tuber number (R = -0.22) and tuber size (R = -0.14), but positively correlated with flowering time (R = 0.091). As with the correlations reported here, prior reports do not show correlations above 0.5, but are highly significant in a much larger population. The genotyping and evaluation of larger breeding populations and additional analytical approaches may offer new insights into the phenotypic impact of DelMut.

### Single gene analysis of predicted deleterious mutations

It is important to mention that mildly DelMut should be analyzed as part of the overall deleterious load of a plant (genetic load). While a single mutation may be of interest and impactful on a single gene basis, it is the combined effect of multiple mildly DelMut that will have larger and discernible phenotypic effect on populations. In our approach, DelMut are predicted based on both their evolutionary conservation and their potential impact on protein sequence and structure. The aggregated and subtle effects of multiple DelMut on proteins function may affect metabolic or physiological processes at varying levels that would later impact the plant performance under certain conditions. As we showed, alleles predicted as highly deleterious on the *PvUI111.03G03490* and *PvUI111.04G120200* genes are in highly conserved protein domains.

The Dihydrolipoyllysine-residue acetyltransferase protein (*PvUI111.03G03490*) forms multimers that are part of the pyruvate dehydrogenase multienzyme complex involved in the super pathway of cytosolic glycolysis in plants, pyruvate dehydrogenase and Tricarboxylic Acid (TCA) cycle (Emes and Tobin, 1993). The amino acid substitution S15L on the protein changes a polar uncharged amino acid (Serine) for a nonpolar and hydrophobic residue (Leucine) which may have implications on the proper protein activity. Another example is the R174G change in the Carboxylesterase 9-related protein (*PvUI111.04G120200*). Arginine is a hydrophobic, charged and polar amino acid that is replaced by a neutral, uncharged and nonpolar glycine. The Carboxylesterase 9-related protein is involved in the methyl indole-3-acetate interconversion in the indole-3-acetic acid (IAA) synthesis. IAA is the most abundant form of auxins, involved in numerous plant growth and development processes and responses to biotic and abiotic stimuli (Woodward and Bartel, 2005). Multiple genes like these, involved in core processes but with sub-optimal functions due to the DelMut may contribute to reduce the overall plant performance.Depending on the context, DelMut may represent an advantage for various purposes. Common bean is known for a higher content of lectins compared to other pulses, which limits its use for animal and human consumption (Bollini and Chrispeels, 1978; Vitale and Bollini, 1995). The selection of DelMut modifying lectin genes could be advantageous. In the parents of the MAGIC population, the gene *PvUI111.04G158600*, a Legume lectin domain protein, had the SNP S04_48751610 [C/T] with a deleterious score of 0.79. Although not at the higher end of the predicted scores, the mutation modifies Alanine 2, a non-polar amino acid for a polar, neutral Threonine (A2T). Similarly, the SNPs S04_48731443 [A/G] (deleterious score 0.59) and S04_48745126 [A/G] (deleterious score 0.53) genotyped in the RILs, accumulated in the lectin genes *PvUI111.04G158400* and *PvUI111.04G158500*, respectively. Depending on the breeding purposes, DelMut predicted with the RF model occurring in genes of interest can be further explored. Lectin could be a model for the analyses of these mutations with deleterious effects based on conservation and protein structure but potentially useful in a practical context.

## Conclusions

In this study, we exploited phylogenetic analyses in legume species to detect selection signatures and predict putatively deleterious mutations in common bean. Middle American beans showed more functional variation (positive selection) in nitrogen-related genes whereas Andean beans in phosphorous-related models. By combining evolutionary and protein information with machine learning, we trained a random forest model to predict DelMut in common bean populations. The model can be implemented in diverse populations with existing genotypic data. Variation in deleterious mutations in the founders and offspring of an artificial MAGIC population of black beans was observed. Based on the high sequencing depth, parents accumulated up to 152 *highly* DelMut (score > 0.85) with burdens up to 17.5K. Under the sequencing conditions and imputation methods, some lines had no predicted *highly* DelMut (score <0.80) highlighting the potential purge of DelMut in populations. We showed *in-silico* how mutations with high deleterious scores (e.g. S04_43764780 [C/G]) are in highly conserved protein domains of core metabolic genes (e.g. involved in TCA cycle). These mutations modify protein sequence with potential impacts on protein function based on amino acid properties changes. Our results contribute to the expanding knowledge of the role of DelMut in plants and breeding populations. Our research also provides novel resources for the acceleration of breeding programs and the development of improved bean varieties. Information on DelMut and genetic load could be incorporated in breeding pipelines aimed at improving selection or prioritizing targets in molecular applications such as multiplex gene editing.

## Data availability statement

All code and data used in this study are available at the McGill University Pulse Breeding and Genetics GitHub page: https://github.com/McGillHaricots/peas-andlove/tree/master/DelMut_genomics

