## Supplementary Figure for "Phylogenetic Analysis and Machine Learning Identify Signatures of Selection and Predict Deleterious Mutations in Common Bean"

**Supplementary material**

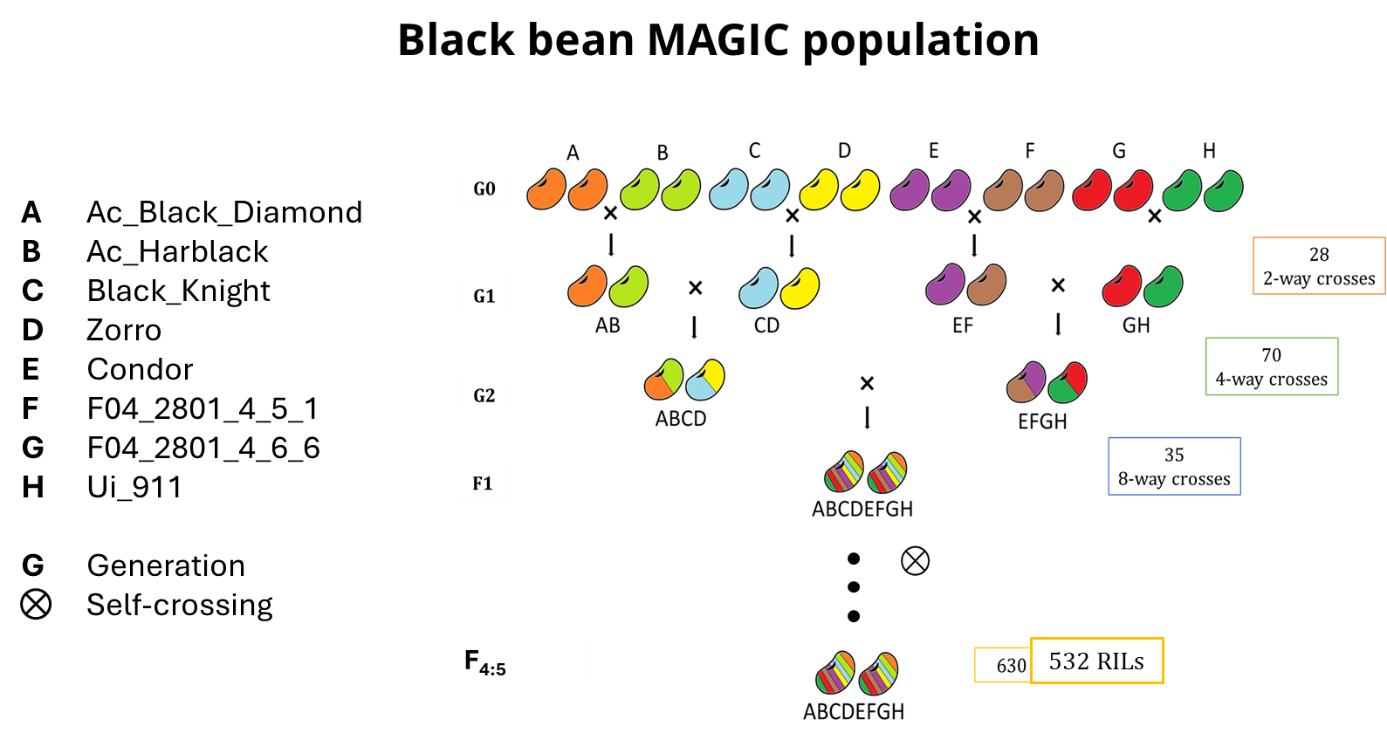

**Figure S1.** Schematic representation of the crossing scheme for the black bean MAGIC population. The two-way crosses were multi-funnel (diallelic) having all the parental combinations. Crosses with the same parents either as female or male were considered reciprocal.

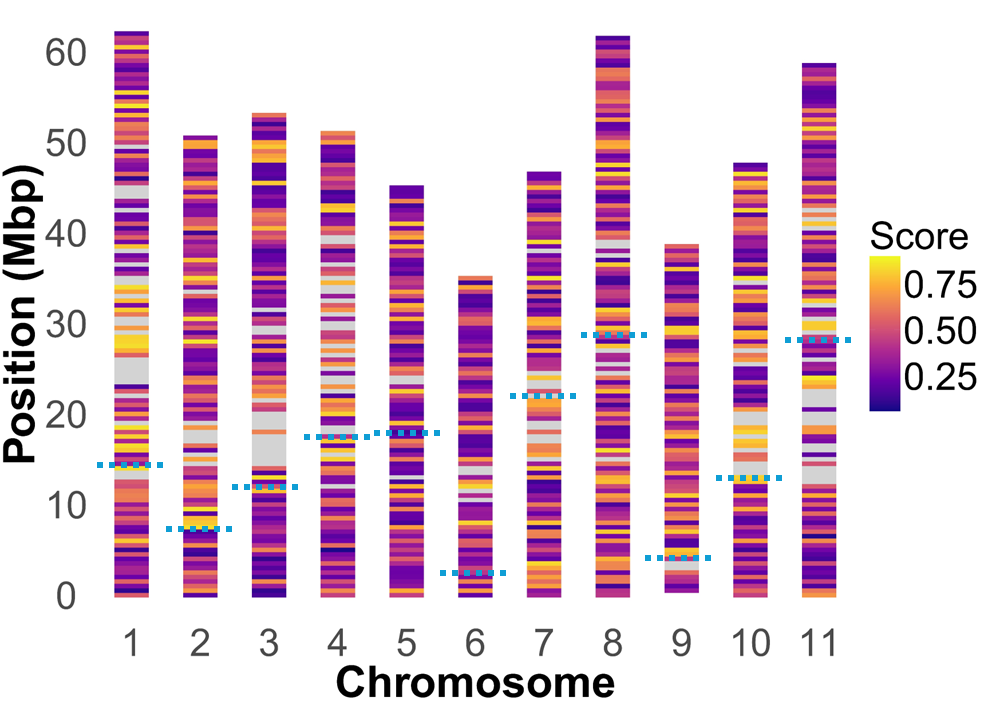

**Figure S2.** Distribution of deleterious scores per chromosome based on the sequencing of the eight *parents* of the black bean MAGIC population.

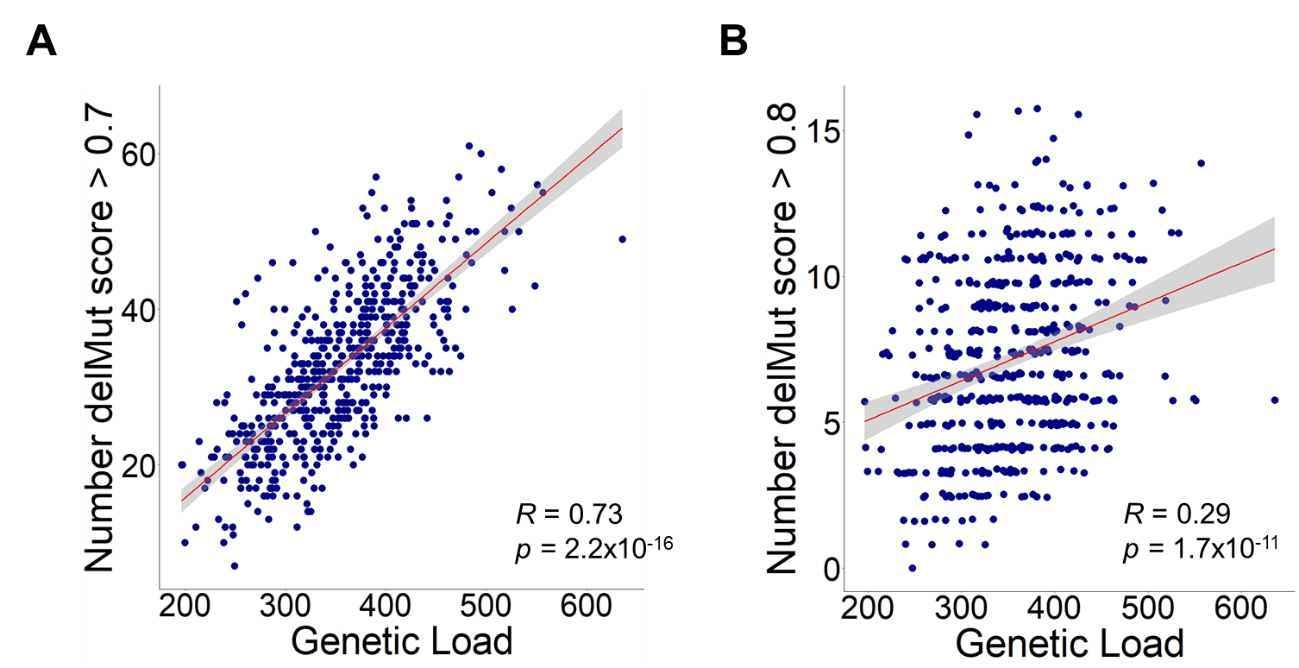

**Figure S3.** Correlation between the number of deleterious alleles with score > 0.70 (A) and > 0.80 (B) and genome-wise genetic load in MAGIC population RILs. Pearson’s correlation coefficient – R is indicated with the p-value of the test.

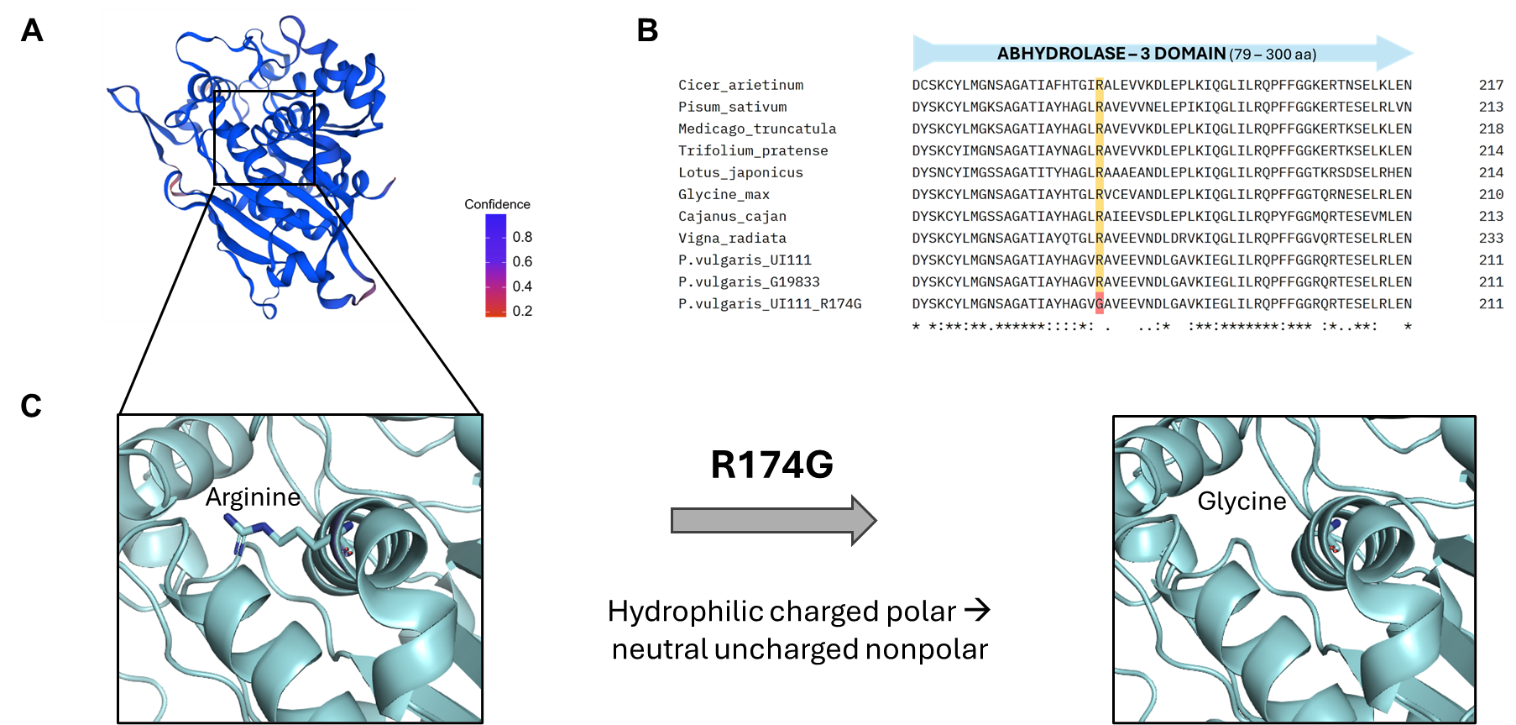

**Figure S4.** Predicted tertiary structure (A) and multiple sequence alignment (B) of the *P. vulgaris* UI111 Carboxylesterase 9-related protein (*PvUI111.04G120200*). Amino acid substitution predicted as deleterious in the AB hydrolase-3 protein domain (C).
